# A novel two-component system controls vancomycin resistance in epidemic *Clostridioides difficile*

**DOI:** 10.1101/2025.08.21.671617

**Authors:** Jessica E. Buddle, Victoria K. Pennington, Joseph A. Kirk, Franziska Faber, Claire E. Turner, Michael A. Brockhurst, Robert P. Fagan

## Abstract

The glycopeptide antibiotic vancomycin is the frontline treatment for *C. difficile* infection in the UK. There have been only sporadic reports of resistance in the clinic but testing is rare so the true resistance landscape is unclear. We have previously shown that resistance can emerge rapidly *in vitro* via distinct but complementary pathways. Strain Bc2, characterised here, is an experimentally evolved derivative of *C. difficile* strain R20291 that displays a 16-fold increase in vancomycin MIC over its parent. Bc2 has point mutations in *dacS*, *bclA3, CDR20291_0794, CDR20291_1871* and *CDR20291_3124 (vnrS)*. By genetically engineering a wild-type vancomycin susceptible strain, we demonstrated that a combination of just two mutations, *dacS*c.798A>T and *vnrS*c.692G>T, both of which encode two-component system histidine kinases, was sufficient to recapitulate Bc2 resistance. We have previously shown that mutations in *dacS* can confer low level resistance via increases in the expression of a D,D-carboxypeptidase DacJ. *dacS*c.798A>T also led to increased transcription of *dacJ* and led to a modest increase in vancomycin MIC. Surprisingly *vnrS*c.692G>T led to overexpression of the *vanG* cluster, which encodes all of the enzymes needed for resistance via substitution of the terminal D-Ala on peptidoglycan lipid II precursors with D-Ser. Genomic analysis of a large collection of European *C. difficile* strains showed that a three gene cluster, which includes *vnrS*, *vnrR* (encoding the cognate response regulator) and an adjacent gene *CDR20291_3123*, is unique to the phylogenetic branch that contains strains belonging to epidemic ribotypes 027 and 176. Transcriptomic analysis of the wider VnrS regulon also revealed an additional previously unknown role in regulating the expression of the flagellum, an important virulence factor in *C. difficile*. Analysis of the response to vancomycin exposure also revealed that *dacJ* is one of a small set of genes that are upregulated shortly after antibiotic stress, even in the absence of mutations that typically lead to its overexpression. Together these data reveal a new synergistic route, needing only two point mutations, by which epidemic lineages of *C. difficile* can attain vancomycin resistance.

**Graphical abstract:** 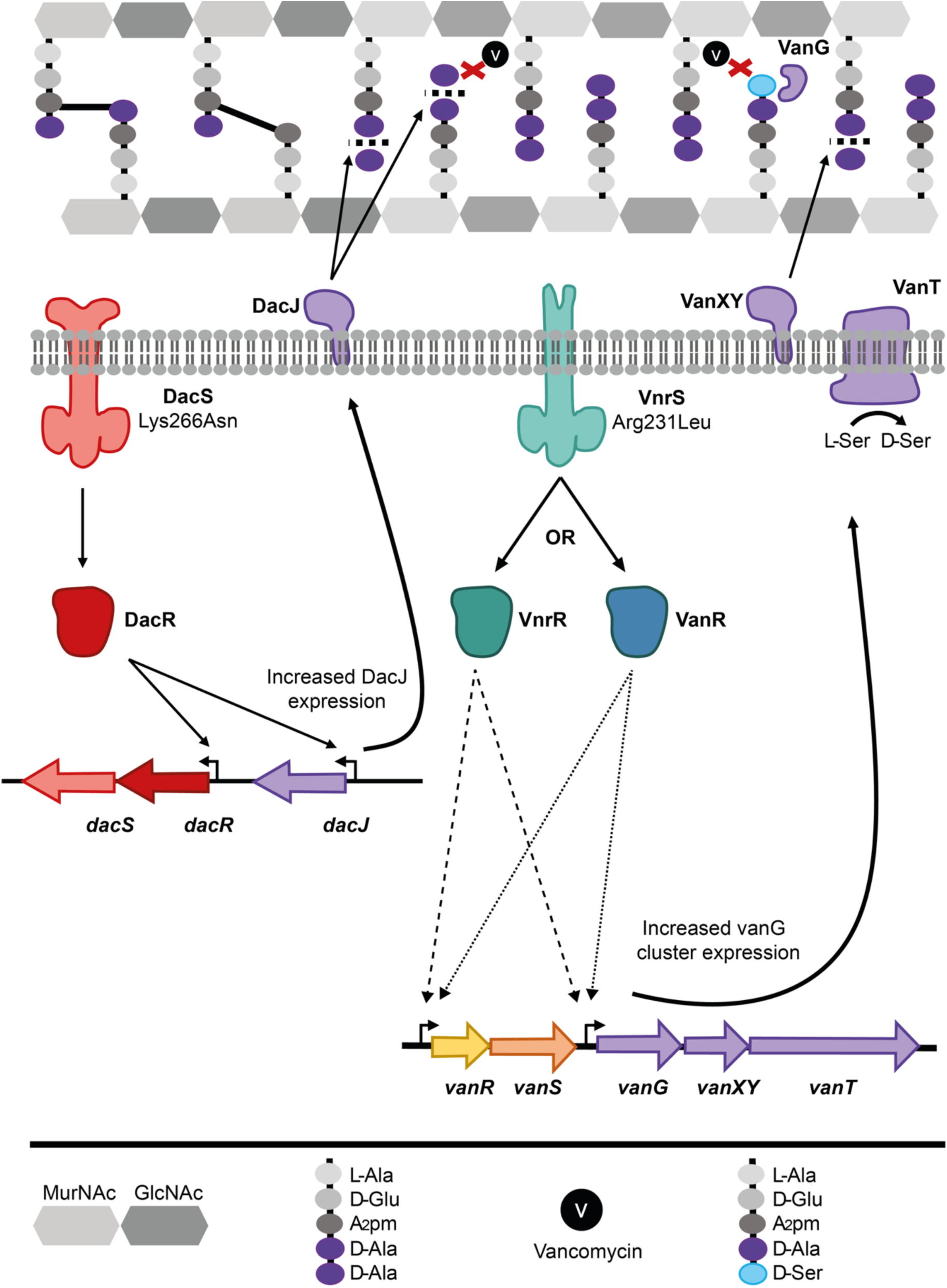

## Introduction

*Clostridioides difficile* is a gram-positive anaerobic bacterium and the leading cause of nosocomial antibiotic-associated diarrhea and colitis worldwide (1). The incidence and severity of *C. difficile* infection (CDI) have risen significantly over the past two decades, posing a substantial burden on healthcare systems and increasing patient morbidity and mortality (2). The primary risk factor for CDI is gut dysbiosis as a result of broad-spectrum antibiotic use, which compromises the colonisation barrier normally provided by a diverse microbiome, allowing *C. difficile* to colonise and proliferate (3). The current front-line treatments for CDI include the antibiotics vancomycin and fidaxomicin, with metronidazole still used in some countries despite increasing resistance (4). However, the use of these agents, particularly vancomycin, can further exacerbate dysbiosis, contributing to disease recurrence in up to 25% of patients (5).

Vancomycin is a glycopeptide antibiotic that inhibits cell wall synthesis by binding to the terminal D-Ala-D-Ala residues of peptidoglycan precursors and sterically hindering crucial transpeptidation and transglycosylation reactions (6). Resistance to vancomycin is well-characterized in several bacterial species, most notably in the Enterococci and *Staphylococcus aureus* (7, 8). In these organisms, resistance is often mediated by proteins encoded within various van gene clusters, that convert the terminal D-Ala-D-Ala to D-Ala-D-Ser or D-Ala-D-Lac that have significantly lower binding affinities for vancomycin (6). While the clinical efficacy of vancomycin against *C. difficile* is clear, indeed it is the drug of choice in many countries (9), concerns are growing due to scattered reports of reduced susceptibility and treatment failure (10, 11). Although a vanG-like gene cluster has been identified in many *C. difficile* strains, this does not appear to confer resistance under normal circumstances (12) and the capacity for and precise mechanisms by which vancomycin resistance may evolve in this pathogen remain largely unknown. Previous work has demonstrated that mutations that impact the expression of vanG cluster genes can lead to limited resistance (13, 14). A small number of additional gene variants have been associated with reduced susceptibility but the mechanistic basis is often unclear (11). We have previously performed a large-scale *in vitro* experimental evolution of vancomycin resistance in the epidemic ribotype 027 *C. difficile* strain R20291 (15), achieving substantial increases in vancomycin MIC in just 250 bacterial generations. We identified over 500 mutations associated with resistance but all resistant strains had mutations affecting two pathways centering around increased expression of the vanG cluster genes or overexpression of a novel D,D-carboxypeptidase DacJ.

Here we mechanistically dissect a vancomycin resistant strain with mutations affecting two distinct two component systems that regulate the expression of DacJ and the genes of the vanG cluster. By recapitulating these mutations in a clean genetic background we reveal that they act synergistically to confer high level resistance. We further characterised the regulon of the vanG-controlling two-component system, which we name VnrRS, and analyse its phylogenetic distribution in a panel of European *C. difficile* clinical isolates.

## Methods

### Strains and growth conditions

*C. difficile* strains were routinely cultured on brain heart infusion (BHI) agar and in TY broth at 37°C in an anaerobic cabinet (Don Whitley Scientific), with an atmosphere composed of 80% N_2_, 10% CO_2_, and 10% H_2_. Where needed culture media was supplemented with thiamphenicol (15 μg/ml; Merck) and colistin (50 μg/ml; Merck) as appropriate. For counter selection against plasmids bearing the *codA* gene, *C. difficil*e defined media with 5-fluorocytosine (CDDM 5-FC) was used as described previously (16). *E. coli* was routinely cultured in Luria–Bertani (LB) broth or agar at 37°C and supplemented with chloramphenicol (15 μg/ml) (Acros Organics) or kanamycin (50 μg/ml) (Merck) as appropriate.

### C. difficile mutagenesis

Mutations were introduced into the R20291Δ*PaLoc* genome using pJAK217, an improved homologous recombination vector generated from our previously published pJAK112 (17). pJAK112 was modified by inverse PCR using oligonucleotides RF1801 and RF1802 (Table S1) to add a well-characterised transcriptional terminator from *Streptococcus pyogenes* (TTATCAACTTGAAAAAGTGGCACCGAGTCGGTGCTTTTTTT (18)) after the *catP* gene to prevent transcriptional readthrough affecting recombination efficiency. Recombination vectors for introduction of *dacS*c.798A>T (pJEB037) and *vnrS*c.692G>T(pJEB038) were generated as previously described (15). Briefly, approximately 2 kb DNA fragments with the mutation of interest in the centre were synthesised by Genewiz (Azenta Life Sciences) and cloned between BamHI and SacI sites in pJAK217. For the reintroduction of the wild-type *vnrS* allele into strain Bc2, mutagenesis plasmid pJEB038 was modified by inverse PCR using oligonucleotides RF2636 and RF2637 to revert the *vnrS*c.692G>T mutation back to the wild-type sequence. The resulting PCR product was ligated to itself, transformed in *E. coli* strain NEB5⍺ and the modified plasmid confirmed by Sanger sequencing of the insert (Azenta Life Sciences). Plasmids were transformed into *E. coli* CA434 and conjugated into *C. difficile* strain R20291Δ*PaLoc* or the evolved vancomycin-resistant derivative Bc2, using standard methods (19). Homologous recombination was carried out as previously described (16) and mutations confirmed by PCR analysis and Sanger sequencing of the altered allele.

### Vancomycin MICs

Vancomycin MICs were determined using standard agar dilution methods (20) as we have described previously (15). Briefly, overnight cultures of *C. difficile* strains in TY broth were diluted to OD_600nm_ 0.1, and 2.5 μl samples were spotted in biological triplicate and technical duplicate onto BHI plates with ranging vancomycin concentrations from 0.25 - 32 µg/ml. MICs were defined as the lowest concentration that did not permit growth of the strain after 48 h incubation.

### RNA purification

RNA was extracted following a hot acid phenol method (21). Triplicate *C. difficile* overnight cultures were subcultured 1:10 in 10 ml TY broth and grown to OD_600nm_ of approx. 0.4 - 0.5. Cells were harvested by centrifugation (4,500 rpm, 4°C, 20 min), resuspended in sodium acetate/SDS buffer (0.1% SDS, 30 mM CH₃COONa, pH 5.5) and transferred to a microfuge tube containing 500 μl of AquaPhenol (Thermo Fisher) which had been heated to 75°C. Samples were incubated at 75°C for 10 min, with regular mixing by inversion, then centrifuged at 21,100 x *g* for 10 min. The supernatant was transferred to a microfuge tube and 500 μl of phenol:chloroform:isoamylalcohol (25:24:1; Thermo Fisher) was added. Samples were centrifuged again as before. The supernatant was transferred to a microfuge tube and 500 μl of isopropanol and 500 μl of sodium acetate (3 M, pH7) was added. After overnight incubation at -20°C, RNA was harvested by centrifugation at 21,100 x g for 30 min at 4°C, washed with 1 ml ice-cold 80% ethanol, and dried by evaporation at room temperature. Precipitated RNA was resuspended in 46 μl of nuclease-free water (Thermo Fisher), residual DNA was removed using the TURBO DNA-free kit (Invitrogen) and RNA yield and quality was assessed by A_260nm_ nano-spectrophotometry.

### qRT-PCR and RNAseq

Single-end 100 bp RNA sequencing was performed by the Core Unit Systems Medicine at the University of Würzburg on a NextSeq 2000 instrument using Truseq adaptors, with the mRNA information in the sense stand. For qRT-PCR cDNA was generated using Superscript III (Invitrogen) following the manufacturer’s protocol. cDNA was quantified by A_260nm_ and concentrations were adjusted to 10 ng/μl. Expression of *dacJ* or *vanG* was measured against an exact copy number control via standardisation with a plasmid containing one copy of each target DNA sequence and normalised against expression of housekeeping gene *rpoA* (21). qRT-PCR target sequences, each ∼200 bp, of genes *dacJ* and *rpoA* (pJEB029) or of *vanG* and *rpoA* (pJEB032) were synthesised and assembled in the pUC-GW-Kan backbone (Genewiz) and were used at a dilution corresponding to a known copy number (2 × 10^8^ per μl). qRT-PCR was performed using SYBR Green JumpStart Taq ReadyMix (Merck) with appropriate forward and reverse primers (Table S1) at concentrations determined by prior optimisation. Templates were cDNA (50 ng in 5 μl), 5 μl of an RT negative control, or 5 μl template plasmid, pJEB029 or pJEB032 serially diluted in lambda DNA (Promega). qRT-PCR was performed in a BioRad CFX Connect Real Time System and cDNA copy numbers were calculated using BioRad CFX manager (v3.1), with data analysed in Microsoft Excel to determine copies per 1,000 copies of *rpoA*. Data were graphed and statistically analysed in GraphPad Prism (v9.0.2).

### Bioinformatics

To assess conservation of the *vnrRS* locus, raw sequence reads from a European collection of *C. difficile* isolates (22)(project accession PRJNA398458) were downloaded and trimmed using Trimmomatic (v0.39, (23)). Trimmed reads were assembled *de novo* using SPAdes (v3.15.5, (24) and the quality of the assemblies were checked using quast. Any that had a total length of <4.0 kb or >4.5 kb were excluded. Assemblies that passed the quality control were annotated using Prokka (v1.14.6, (25) and pangenome analysis was performed using panaroo (v1.14.6, (26)). From this, the presence of *CDR20291_3123-3125* genes was determined and, where present, the sequences of these genes were compared. Gene presence/absence was related back to the previously determined ribotype for each isolate (22). A phylogenetic tree was produced from the panaroo-generated core gene alignment using RAxML (27).

Differential expression analysis was performed on sequencing data provided by the University of Würzburg Core Unit Systems Medicine facility. Briefly, raw fastq reads were trimmed using Trimmomatic (v0.30, (23)) with a sliding window quality cutoff of Q15, to remove adapters and low quality reads. Trimmed reads were checked using FastQC (v0.11.9, (28)) to ensure sufficient quality for analysis. Reads were aligned to the *C. difficile* R20291 reference genome (accession number: FN545816) using BWA-mem (v0.7.17, (29)), and sorted using SAMtools (v1.43, (30)). The resulting bam files were then used in conjunction with the *C. difficile* R20291 feature file (GFF) to generate count files, using HTSeq-count (v2.0.5, (31)). The count files, indicating the number of reads mapping to a particular feature, were then compiled into a count matrix, which was used as the input for differential expression (DE) analysis. Differential expression was performed using the DESeq2 package (v3.21, (32)) in RStudio (v4.1.0), using a negative binomial generalized linear model, accounting for biological variation among replicates. Genes were considered significantly differentially expressed if the Benjamini-Hochberg FDR adjusted p-value (padj) was ≤ 0.01, and the log₂ fold change was greater than 2 or less than -2. Differential expression across the genome was visualised in RStudio (v4.1.0) using gene loci as a proxy for genomic position.

## Results

### Recapitulation of mutations in two histidine kinases is sufficient to confer vancomycin resistance

Previous hybrid sequencing of evolved vancomycin resistant strain Bc2 (15) identified 5 mutations not seen in the parental or control evolved strains: *dacS*c.798A>T, *bclA3*c.640_642delATAinsGCAGAT, *CDR20291_0794*c.732delA, *CDR20291_1871*c.2178delA and *CDR20291_3124*c.692G>T (hereafter *vnrS* for van-related sensor). Three of these encode proteins with no plausible connection to vancomycin resistance: *bclA3* encodes a putative exosporium glycoprotein with no known function in the vegetative cell, *CDR20291_0794* encodes a conserved hypothetical protein and *CDR20291_1871* encodes a putative ABC transporter permease. Initial efforts to recapitulate resistance focussed on two genes encoding two-component system histidine kinases *dacS* and *vnrS*. We have previously shown that DacS controls expression of a D,D-carboxypeptidase DacJ that can confer vancomycin resistance when overexpressed. The gene adjacent to *vnrS*, *CDR20291_3125* (hereafter *vnrR*, van-related regulator), encodes the likely cognate response regulator but the regulon of this two-component system has yet to be characterised. To determine the contribution of the *dacS* and *vnrS* mutations to vancomycin resistance in Bc2, each mutation was recapitulated either alone or in combination in the parental strain R20291Δ*PaLoc* (Table 1). *dacS*c.798A>T alone resulted in only a 2-fold increase in vancomycin MIC and *vnrS*c.692G>T 4-fold. The combination of the two mutations, however, fully recapitulated the 16-fold increased MIC seen in the evolved Bc2, suggesting that these mutations act synergistically to confer high-level resistance. To confirm the contribution of *vnrS* we performed the analogous experiment in evolved strain Bc2, reverting the *vnrS*c.692G>T allele back to the wild-type sequence. This mutation reduced the Bc2 MIC from 16 µg/ml to 2 µg/ml, confirming both the contribution of VnrS and the synergistic interaction with *dacS*c.798A>T.

**Table 1.** Vancomycin MICs of strains recapitulating Bc2 mutations.

| Strain | MIC (µg/mL) |
| --- | --- |
| R20291Δ*PaLoc* | 1 |
| Bc2 | 16 |
| R20291Δ*PaLoc* *dacSc.798A>T* | 2 |
| R20291Δ*PaLoc* *vnrSc.692G>T* | 4 |
| R20291Δ*PaLoc* *dacSc.798A>T vnrSc.692G>T* | 16 |
| RBc2 *vnrSc.692T>G* | 2 |

### VnrS is a novel regulator of the *vanG* cluster

We have previously shown that the DacRS two-component system regulates its own expression and that of DacJ, a D,D-carboxypeptidase, and that amino acid substitutions in DacS (Glu238Asp or Val183Ala) that result in increased transcription of *dacJ* confer vancomycin resistance (15). Given the proximity of the DacS mutation in Bc2 (Lys266Asn) to one we have previously linked to vancomycin resistance (Glu238Asp), we hypothesised that Bc2 would also overexpress DacJ. To assess this, we used qRT-PCR to quantify *dacJ* mRNA in our panel of strains, including the parental strain R20291Δ*PaLoc*, evolved Bc2 and engineered mutants in *dacS* and *vnrS* in all combinations (Fig. 1A). All strains with the *dacS*c.798A>T mutation displayed increased *dacJ* transcription, consistent with this being the mechanism of decreased vancomycin susceptibility observed. In all cases, the *vnrS*c.692G>T had no significant effect on *dacJ* expression suggesting that this mutation is influencing vancomycin resistance via a distinct but complementary mechanism. As we have previously observed synergistic effects between the DacJRS and vanG pathways, we assessed the expression of *vanG*, the first gene in the *vanGXYT* operon (Fig. 1B). The *dacS*c.798A>T mutation had no apparent effect on *vanG* expression but surprisingly, all strains with the *vnrS*c.692G>T mutations displayed increased transcription of *vanG* and reversion of the mutation in Bc2 reduced the *vanG* transcript back to wild-type levels. From this we conclude that the VnrRS two-component system is a previously unknown regulator of the *vanG* cluster, a locus encoding the regulatory two-component system VanRS and all of the enzymes necessary for substitution of the terminal D-Ala-D-Ala on peptidoglycan precursors for D-Ala-D-Ser, VanG, VanXY and VanT (33).

**Figure 1.**
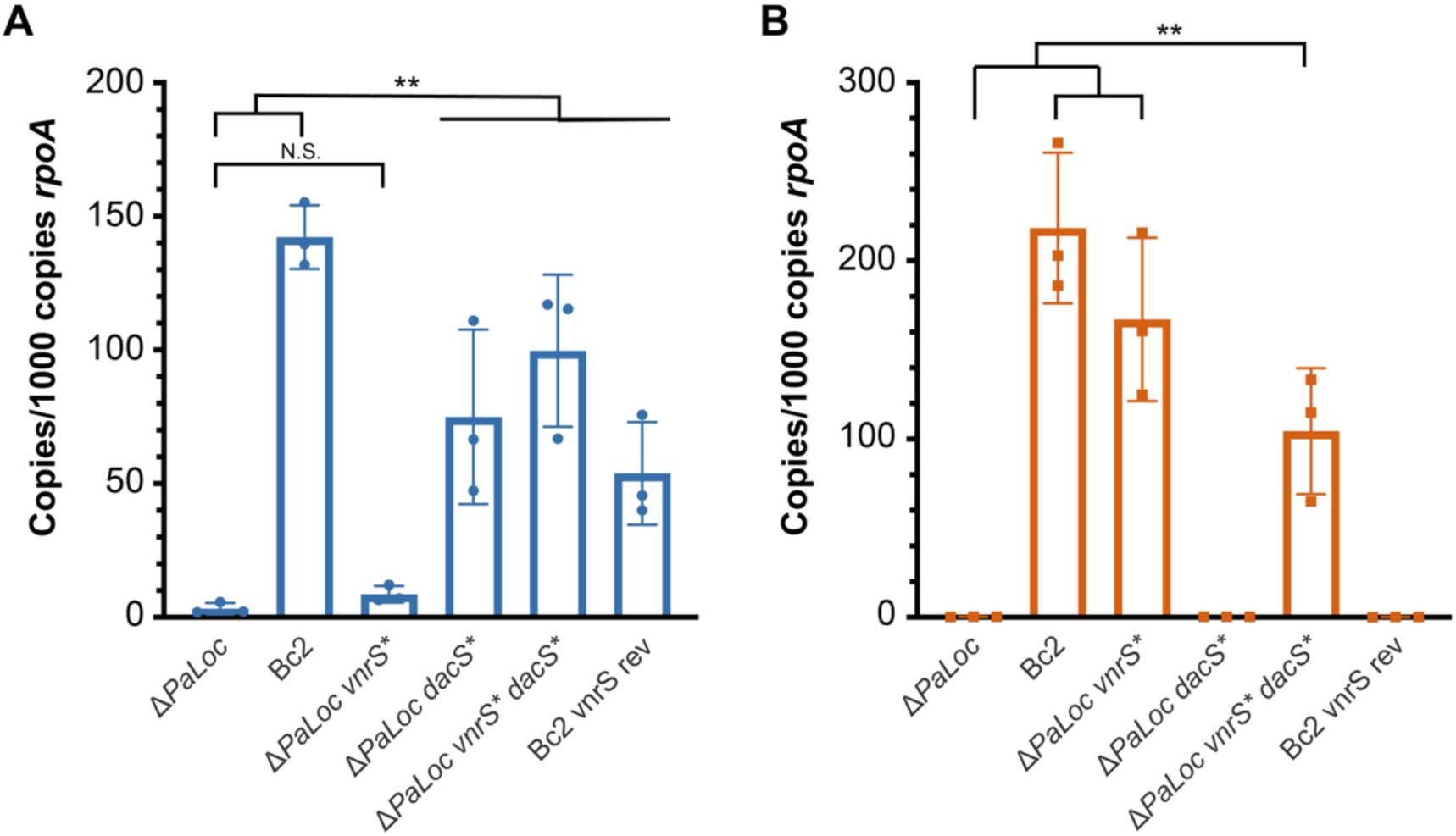
Resistance conferring mutations lead to overexpression of *dacJ* and *vanG*. qRT-PCR analysis of *dacJ* (**A**) and *vanG* (**B**) expression in R20291ΔPaLoc, Bc2, R20291ΔPaLoc *vnrS*c.692G>T (*vnrS*\*), R20291ΔPaLoc *dacS*c.798A>T (*dacS*\*), and R20291ΔPaLoc *vnrS*c.692G>T *dacS*c.798A>T or Bc2 with *vnrS* reverted back to the wild-type sequence (*vnrS* rev). Expression was quantified against a standard curve and normalised relative to the house-keeping gene *rpoA*. Assays were performed in biological and technical triplicate. Shown are the means and standard deviations (bars) with the means of individual biological replicates overlaid as a scatter plot. Statistical significance was calculated using a two-way ANOVA with the Tukey–Kramer test, ** = P < 0.001

### *vnrRS* appears unique to epidemic *C. difficile* strains

We noted that the common laboratory strain 630 lacked a clear homologue of *vnrS*. Closer examination of the equivalent genomic region (*CD630_32640-32650*) revealed that a three-gene cluster (equivalent to *CDR20291_3123-3125*) is entirely absent in strain 630 (Fig. 2A). In addition to the two-component system encoded by *vnrRS* (*CDR20291_3124-3125*), the adjacent *CDR20291_3123* encodes a putative ABC transporter ATP-binding protein. Expanding genomic analysis to a collection of 552 publicly available European *C. difficile* genomes (22), representing 10 different ribotypes, revealed that only isolates belonging to the closely related ribotypes 027 and 176 have the three-gene cluster that includes *vnrS* (Fig. 2B) and the remainder have the same (or similar) genomic organisation as seen in strain 630. All ribotype 027 and 176 strains analysed possessed the *CDR20291_3123-3125* cluster and in each case the amino acid sequence of the encoded proteins was identical.

**Figure 2.**
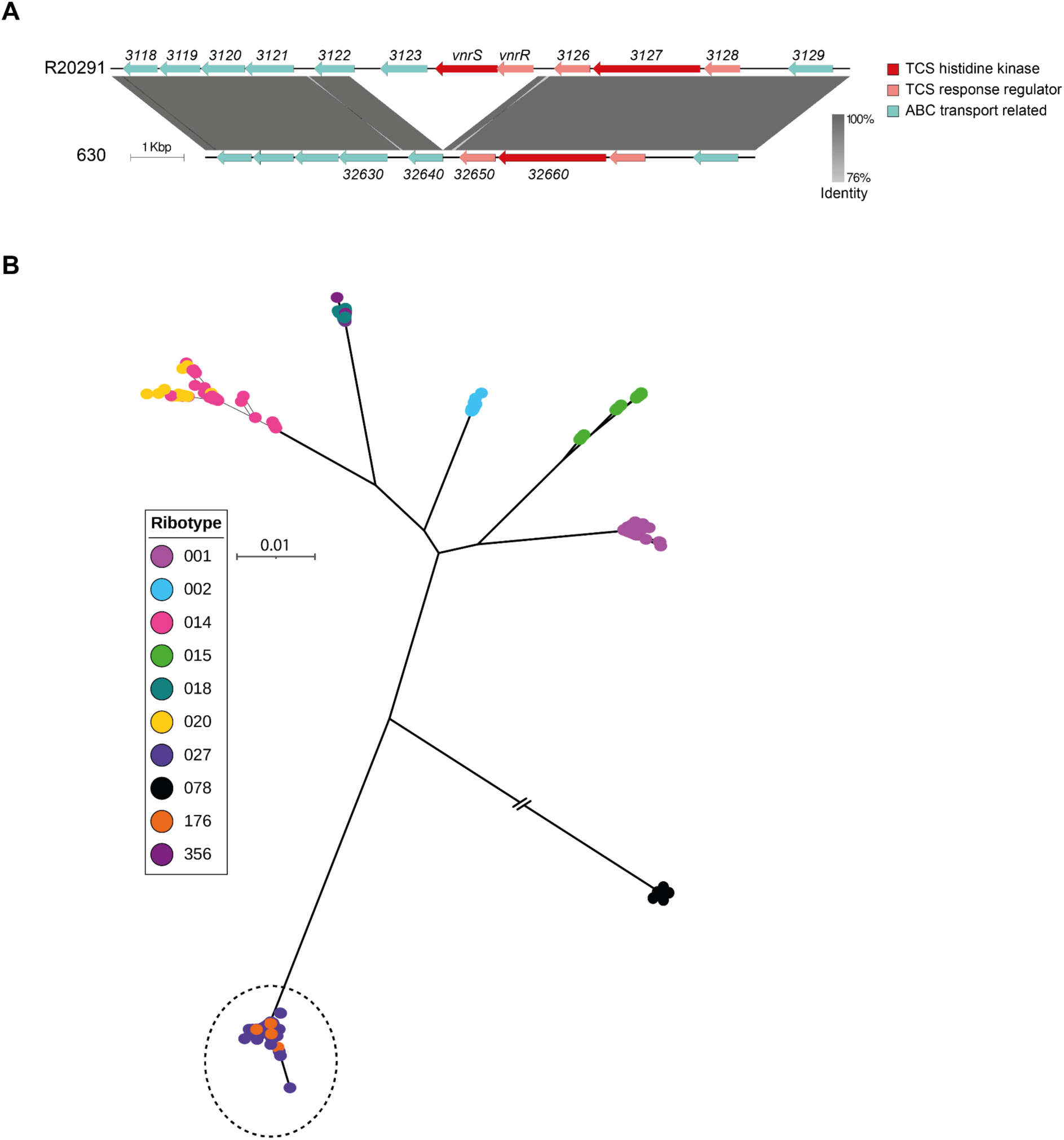
Phylogenetic distribution of *vnrRS*. **A** Comparison of the *vnrRS* locus in *C. difficile* R20291 with the equivalent region in the genome of strain 630. Visualisation generated with Easyfig (34). Genes encoding putative ABC transporter components are shown in blue with two-component system histidine kinases and response regulators in red and pink respectively. **B** Maximum likelihood phylogenetic tree based on a core genome alignment of 552 *C. difficile* genomes from a European epidemiological surveillance dataset (22). Only strains belonging to ribotypes 176 and 027 (circled) possess the three gene cluster that includes *vnrRS* and the adjacent ABC transporter ATP-binding protein-encoding gene. The ribotype 078 lineage branch is shortened for illustrative purposes. The scale bar represents the number of substitutions per site.

### The VnrRS regulon includes the flagellum

To define the wider VnrRS regulon, a clean *vnrS* deletion (R20291Δ*PaLoc*Δ*vnrS*) was constructed and RNAseq was performed on exponentially growing cultures to identify genes impacted by the deletion. Overall, 40 genes showed differential expression (Supp. Data S1, Fig. 3A). Loss of *vnrS* led to significant upregulation of 11 genes, including the downstream *CDR20291_3123*, which encodes a putative ABC transporter component. Interestingly, *CDR20291_3126*, *CDR20291_3127* and *CDR20291_3128,* just upstream of *vnrS,* which encode an additional histidine kinase and two two-component response regulators, also showed significant upregulation in the absence of *vnrS*. Although the regulons of these two-component systems are not known, repression of these genes by VnrRS likely has widespread indirect effects. Both *argJ* and *argF*, encoding members of the arginine biosynthesis pathway, also showed significant upregulation in the absence of *vnrS*, suggesting a role for VnrRS in the modulation of arginine metabolism.

**Figure 3.**
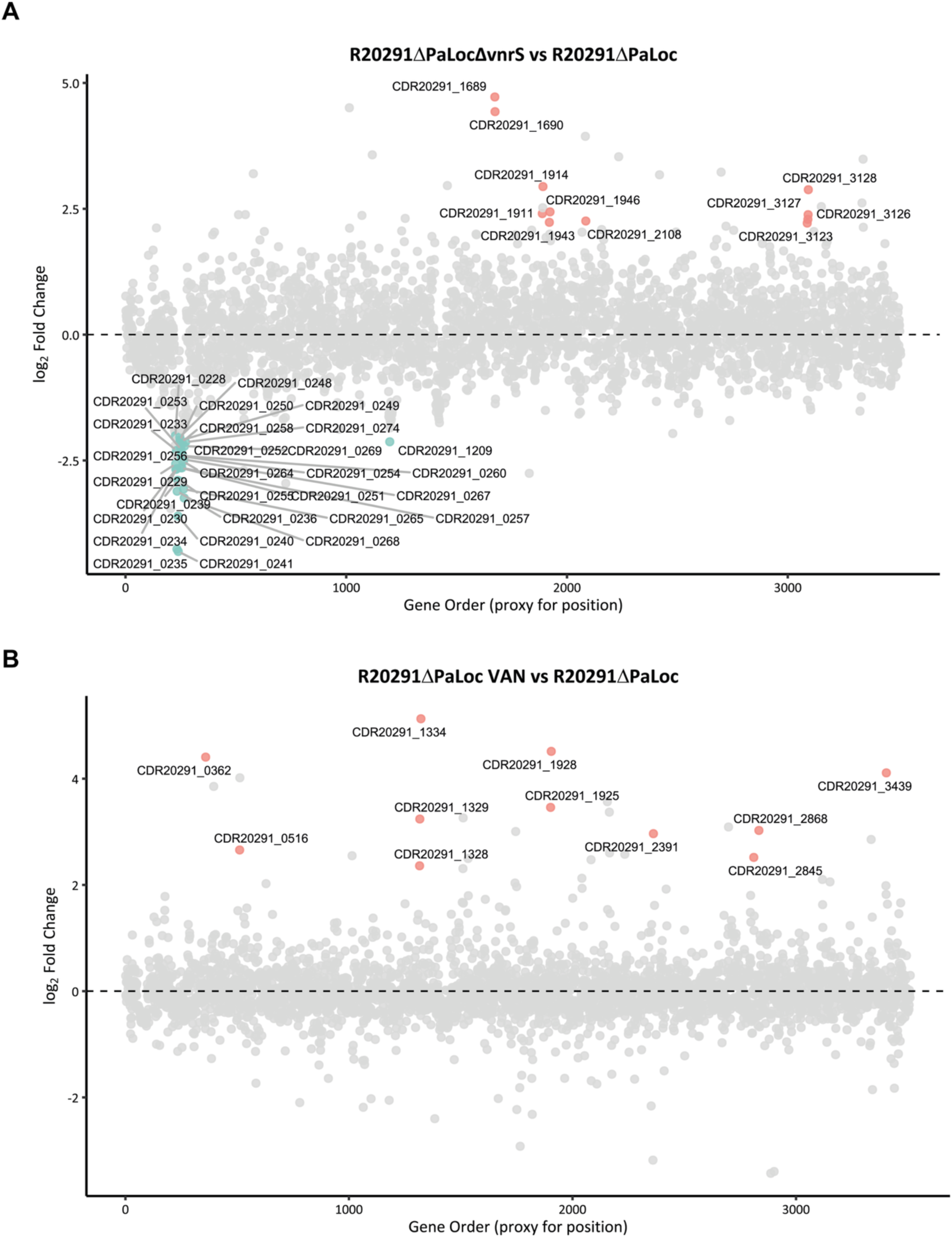
A. Log₂ fold change of every gene across the R20291Δ*PaLoc*Δ*vnrS* genome compared to R20291Δ*PaLoc*. Significantly differentially expressed genes coloured pink (upregulated in R20291Δ*PaLoc*Δ*vnrS*) or blue (downregulated in R20291Δ*PaLoc*Δ*vnrS*) are labelled with locus tags. Locus tag number was used as a proxy for genome position. Differential expression was considered significant if padj was ≤ 0.01, and log₂ fold change was greater than 2 or less than -2. **B** Log₂ fold change of every gene across the R20291Δ*PaLoc* genome during 0.5x MIC vancomycin challenge compared to normal conditions. Significantly differentially expressed genes coloured pink (upregulated in presence of 0.5x MIC vancomycin).

Strikingly, of the 29 genes downregulated in the absence of *vnrS*, 28 are associated with flagellar biosynthesis – spanning biosynthetic proteins, core structural components, assembly and export machinery and regulatory proteins, suggesting VnrRS is an important positive regulator of flagella expression. Among the downregulated genes were *CDR20291_0240* (*fliC*), encoding the major flagellin subunit, *CDR20291_0265* (*fliQ*), encoding flagellar export protein, and *CDR20291_0269* (*fleN*), encoding the flagellar number regulator. The genes displaying the highest levels of differential expression were *CDR20291_0241*, encoding a putative glycosyltransferase, and *CDR20291_0235,* predicted to encode a carbon storage regulator, both located in a locus implicated in glycosylation of the flagella (35).

### DacJ expression is upregulated in response to vancomycin

To understand the early transcriptional responses which occur upon vancomycin exposure, R20291Δ*PaLoc* was challenged for 25 minutes during exponential growth with 0.5x MIC vancomycin (0.5 µg/mL). RNAseq and differential expression analysis was performed as above, to identify genes modulated in the early stages of vancomycin exposure (Fig. 3B). No significantly downregulated genes were identified. Among the 11 significantly upregulated genes are two involved in oxidative stress responses, *CDR20291_2868* (encoding thioredoxin) and *CDR20291_1925* (encoding flavodoxin), consistent with vancomycin exposure generating oxidative stress (36). *CDR20291_0362*, encoding an asparagine synthetase, also showed significant upregulation. We previously identified a mutation in this gene (*CDR20291_0362*c.341T>A) in an evolved vancomycin resistant strain (15) and the encoded protein AsnB1 has been implicated in peptidoglycan amidation in the presence of vancomycin (37). Strikingly, one of the most upregulated genes, with a log_2_ fold change of over 4, was *CDR20291_3439*, encoding the D,D-carboxypeptidase DacJ. We have shown that overexpression of DacJ reduces vancomycin sensitivity in multiple laboratory evolved isolates (15). Upregulation of this gene during vancomycin challenge suggests that even in the absence of mutations conferring DacJ overexpression, DacJ is involved in early responses to vancomycin.

## Discussion

We have previously shown that vancomycin resistance can evolve rapidly in *C. difficile* and identified a large number of mutations associated with resistance *in vitro* (15). One strain isolated during that experimental evolution, Bc2, was found to have mutations in *dacS*, *bclA*, *CDR20291_0794*, *CDR20291_1871* and *vnrS*. Here we have shown that just two of these mutations, *dacS*c.798A>T and *vnrS*c.692G>T, are sufficient to confer resistance. *dacS*c.798A>T (encoding two-component system histidine kinase DacS Lys266Asn) leads to increased expression of DacJ, a D,D-carboxypeptidase that we have previously linked to vancomycin resistance (15). DacJ D,D-carboxypeptidase activity reduces the prevalence of vancomycin binding sites in the cell wall, likely reflecting a conversion of typical pentapeptide lipid II peptidoglycan precursors to tetrapeptides, lacking the terminal fifth position D-Ala residue. This can be tolerated in *C. difficile* due to the overwhelming reliance on 3-3 crosslinks in the cell wall (38), formation of which does not require a pentapeptide substrate (39). The two-component system encoded by *vnrRS* has not been characterised previously and, surprisingly, we found that the *vnrS*c.692G>T mutation resulted in overexpression of *vanG*, a well-characterised mechanism by which *C. difficile* can acquire vancomycin resistance (13, 15). Although it has not been experimentally confirmed in *C. difficile*, the vanG cluster encodes all of the enzymatic activities needed for substitution of the terminal D-Ala-D-Ala on lipid II precursors for D-Ala-D-Ser which has a significantly lower affinity for vancomycin (6). Interestingly, the genes of the vanG cluster were not identified among the normal VnrRS regulon suggesting that the upregulation of *vanG* seen in strains carrying *vnrS*c.692G>T is a direct consequence of this mutation rather than a simple augmentation of normal VnrS function. Instead, it seems that the house keeping function of VnrRS is a previously unexpected role in regulation of flagella biosynthesis. One intriguing possibility is that the amino acid change we observed in VnrS (Arg231Leu) results in phosphotransfer promiscuity that allows crosstalk with the VanR response regulator that typically controls expression of the vanG cluster. Crosstalk affecting the vanG cluster would not be unprecedented (40) and it has been shown that mutations in the Dimerization and Histidine Phosphotransferase (DHp) domain of a histidine kinase can alter the fidelity of HK-RR communication (41) However, characterisation of the mechanistic basis of VnrRS control of vanG will require further analysis.

*vnrRS* is not part of the core *C. difficile* genome and instead appears to be limited to certain lineages, in the 552 strain European clinical dataset (22) analysed here it was only found in the closely related ribotype 027 and 176 strains. As a result, these clinically important strains have an additional route to reduced vancomycin susceptibility, requiring just a single point mutation to achieve first step resistance.

We have previously shown that emerging vancomycin resistance is frequently accompanied by fitness defects that would likely impact the success of the resistant strain in natural infection (15). However, it is likely that additional compensatory mutations that ameliorate the fitness impacts of resistance conferring mutations will evolve in time. It is critical that we fully understand the capacity for and mechanisms by which *C. difficile* may evolve resistance to vancomycin to guide infection control and prescription practices.

## Author statements

### Author contributions

Conceptualisation: Jessica E. Buddle, Michael A. Brockhurst, Robert P. Fagan. Data curation: Jessica E. Buddle.

Formal analysis: Jessica E. Buddle, Claire E. Turner, Robert P. Fagan. Funding acquisition: Michael A. Brockhurst, Robert P. Fagan.

Investigation: Jessica E. Buddle, Victoria K. Pennington, Joseph A. Kirk.

Methodology: Jessica E. Buddle, Victoria K. Pennington, Joseph A. Kirk, Franziska Faber, Claire E. Turner, Michael A. Brockhurst, Robert P. Fagan.

Project administration: Robert P. Fagan.

Supervision: Claire E. Turner, Michael A. Brockhurst, Robert P. Fagan. Visualisation: Jessica E. Buddle, Claire E. Turner, Robert P. Fagan

Writing - original draft: Jessica E. Buddle, Claire E. Turner, Michael A. Brockhurst, Robert P. Fagan.

Writing - review and editing: All authors.

---

**Table S1.**
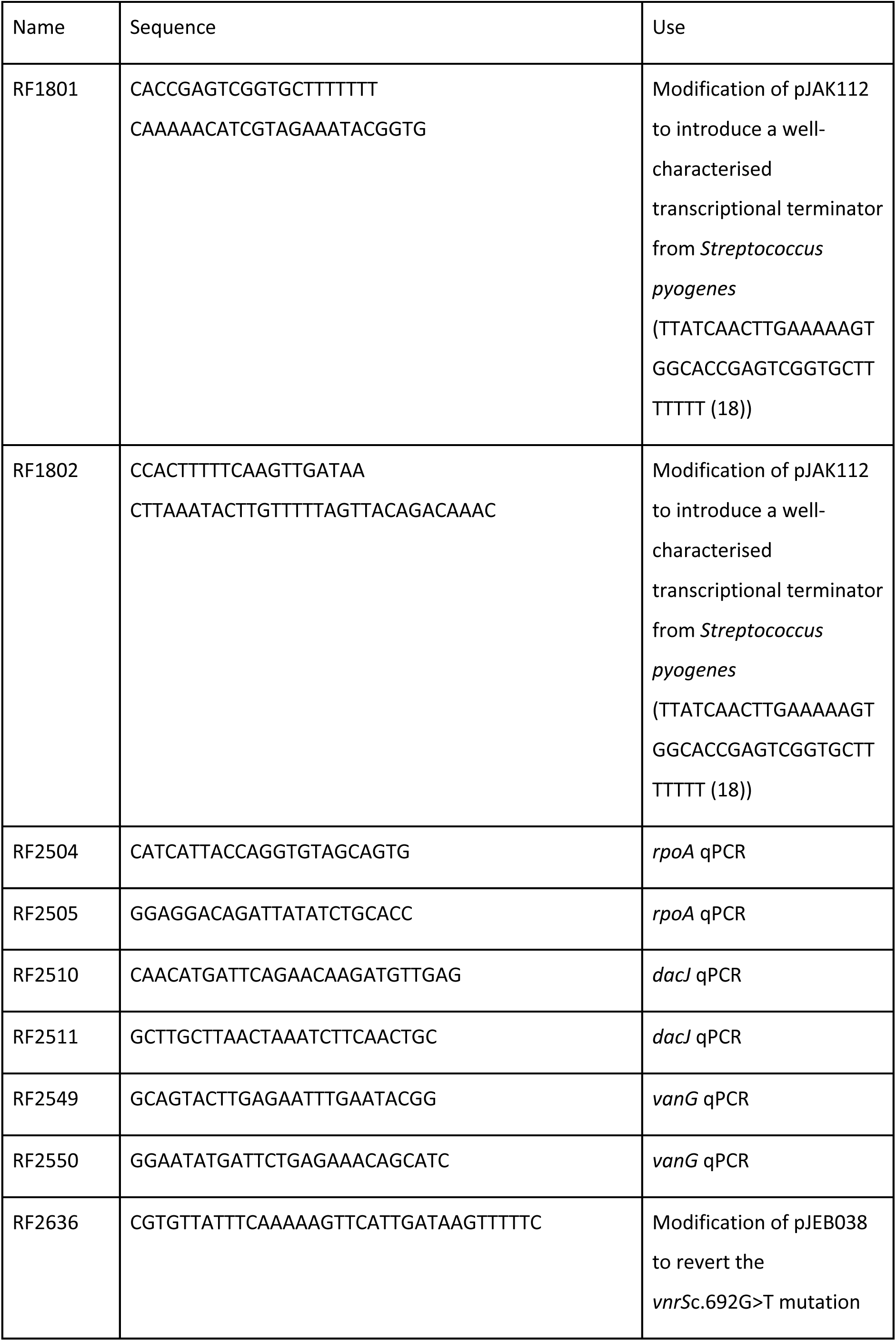
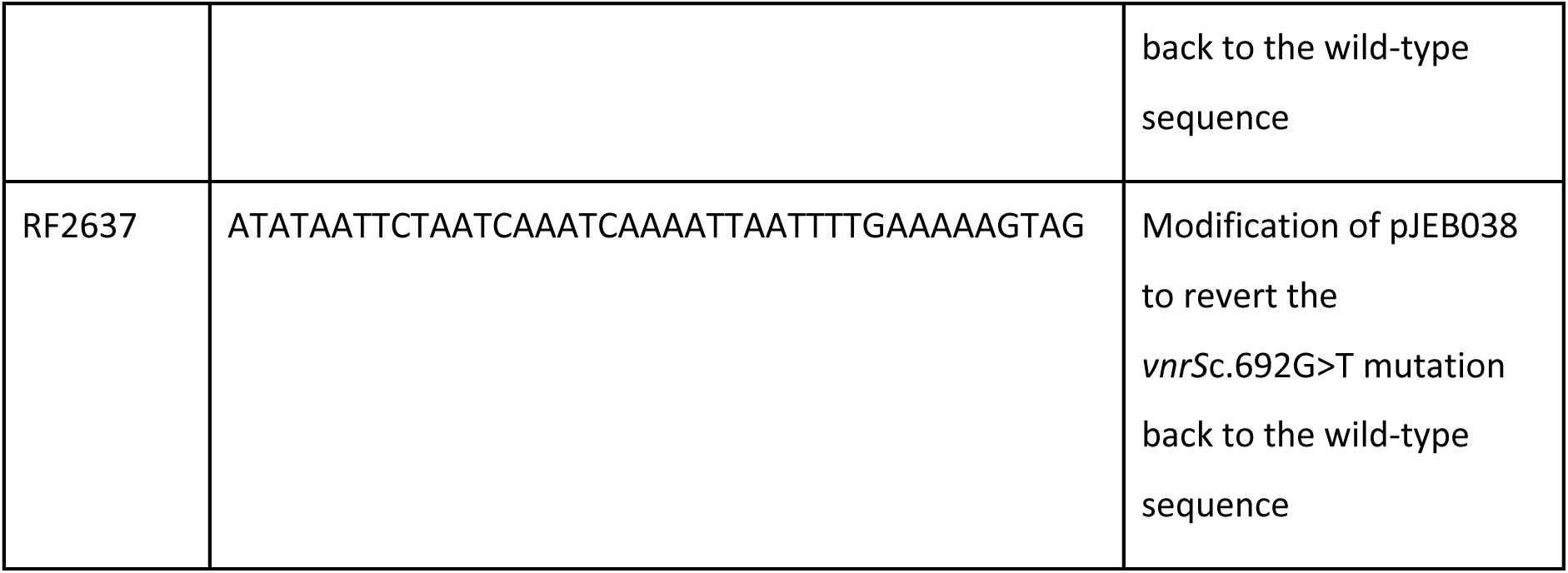
Oligonucleotides used in this study.

